# Residence Time Analysis of RNA Polymerase Transcription Dynamics: A Bayesian Sticky HMM Approach

**DOI:** 10.1101/2020.07.28.132373

**Authors:** Zeliha Kilic, Ioannis Sgouralis, Steve Pressé

## Abstract

The time spent by a single RNA polymerase (RNAP) at specific locations along the DNA, termed “residence time”, reports on the initiation, elongation and termination stages of transcription. At the single molecule level, this information can be obtained from dual ultra-stable optical trapping experiments, revealing a transcriptional elongation of RNAP interspersed with residence times of variable duration. Successfully discriminating between long and short residence times was used by previous approaches to learn about RNAP’s transcription elongation dynamics. Here, we propose an approach based on the Bayesian sticky hidden Markov models that treats all residence times, for an E. Coli RNAP, on an equal footing without a priori discriminating between long and short residence times. In addition, our method has two additional advantages, we provide: full distributions around key point statistics; and directly treat the sequence-dependence of RNAP’s elongation rate.

By applying our approach to experimental data, we find: no emergent separation between long and short residence times warranted by the data; force dependent average residence time transcription elongation dynamics; limited effects of GreB on average backtracking durations and counts; and a slight drop in the average residence time as a function of applied force in RNaseA’s presence.

**STATEMENT OF SIGNIFICANCE:** Much of what we know about RNA Polymerase, and its associated transcription factors, relies on successfully discriminating between what are believed to be short and long residence times in the data. This is achieved by applying pause-detection algorithms to trace analysis. Here we propose a new method relying on Bayesian sticky hidden Markov models to interpret time traces provided by dual optical trapping experiments associated with transcription elongation of RNAP. Our method does not discriminate between short and long residence times from the offset in the analysis. It allows for DNA site-dependent transition probabilities of RNAP to neighboring sites (thereby accounting for chemical variability in site to site transitions) and does not demand any time trace pre-processing (such as denoising).

## INTRODUCTION

The last two decades have seen a surge in single molecule experiments that probe transcriptional dynamics by a number of methods (1–19) including optical force trapping (12, 20–29).

Optical force trapping of single molecules has provided a unique window into the transcriptional dynamics of individual RNAP molecules (30). However, insight drawn from such experiments are highly sensitive to analysis methods of the time traces generated by optical force trapping.

Insights on RNAP transcription include (25, 30, 31): 1) the sequence-dependence of long residence time events (21, 22, 24, 25); 2) the heterogeneity in residence time statistics across RNAP molecules (21, 22, 25); 3) the role of transcription elongation factors such as GreA, GreB, NusG on residence time (20, 22); 4) the role of interactions between the nascent RNA chain and the RNAP molecule and its possible role in the subtle change in residence time statistics as a function of transcriptional duration (20–22, 24, 25); and 5) the possibility of off RNAP pathways resulting in long (trapped) residence times (20–22, 24).

These insights are drawn from time series analysis. The analysis is inherently difficult because of intrinsic noise in time traces obtained from optical trapping arising mainly from thermal fluctuations (30) and extrinsic noise arising mainly from measurement errors (32–35). To facilitate the analysis, experimental time traces are often treated using pause-detection algorithms (20–25) to draw insight on RNAP’s transcriptional residence time dynamics (20, 21, 23–25). These algorithms involve extensive pre-processing such as downsampling and denoising of the data. Subsequently, the user must pre-specify a duration threshold to discriminate between different types of residence time (such as short residences and longer residences) (20, 21, 23, 25). As it is well known, such methods may eliminate fast transitions and introduce artificial states in the experimental time traces (34, 36–39). What is more, in these two-stage methods: 1) the denoising is not achieved simultaneously, and thus self-consistently, alongside the determination of the residence time kinetics; and 2) the physics dictating the noise’s origin, and thus the nature of the noise statistics, is not exploited in the analysis. Thus noise fluctuations anticipated by physical considerations may be misinterpreted as kinetic transitions of the underlying molecule and affect residence time duration estimates. These reasons are chiefly why hidden Markov models (HMMs) have become the workhorse of Biophysics time series analysis in the first place, for example in smFRET (35, 39) and in force spectroscopy (25, 32–34). HMMs have been used in the analysis of optical trapping experiments as they pertain to transcription residence time dynamics; however, in the application of plain HMMs thresholds are often still invoked to discriminate long from short pauses (20–22, 24, 25, 40). Furthermore, plain HMMs assume that the kinetics of RNAP remain homogeneous in space (36–39). A variant of the plain HMM was proposed to find the step distribution of RNAP based on the prescribed residence time distributions (uniform, gamma or exponential distributions) from the analysis of optical trapping experiments (40). This proposed method requires data downsampling and also requires the data to be split into smaller segments for faster convergence of the method (40). Additionally, implementing the method in (40) requires that backtracked motion be removed from the analyzed data for applied forces above a threshold 15 pN due to rarely occuring backtracking events. Similarly, estimating the step distribution of RNAP based on *maximum likelihood* approaches was previously carried out for magnetic tweezer experiments (41, 42). This *maximum likelihood approach* also requires pre-processing of the acquired data from magnetic tweezer experiments. Furthermore, this method appearing in (41, 42) is limited to point estimates for the step distribution rather than providing full step distributions. Recently, a slightly different optimization approach was presented in (24) for the data analysis of optical trapping experiments. This approach is based on estimating the crossing times of a set of nucleotides (pausing sites) on the transcribed DNA template. While this new method in (24) is not a pause detection algorithm, it relies on investigating residence time distribution of RNAP based on crossing time distributions. This method in (24) is still subject to heavy data pre-processing prior to the analysis.

What we need instead is a method of analysis that has the following features: 1) can achieve everything the HMM already can (namely exploit noise statistics rigorously in learning transition probabilities); 2) captures the inhomogeneity in space of the transition kinetics (i.e., transition probabilities) of RNAP from base pair (bp) to bp as it evolves in time and avoids thresholds to separate short from long residence time in the analysis; 3) provides not only error bars but also full distributions over all point statistics (such as average residence time or average backtrack durations).

Here, we propose a novel method that satisfies all three criteria above for the analysis of time traces generated by single molecule dual optical trap experiment as illustrated in Figure 1. We leverage the strengths of HMMs to capture spatially inhomogeneous kinetics. Additionally, as transition probabilities are anticipated to be heterogeneous in space, we consider a generalization of the HMM termed the sticky HMM (43, 44) that achieves point 2 above. This allows to avoid over-interpreting noise in the experimental data from the inherently heterogeneous kinetics where transition probabilities vary depending on the transcribed bps. Finally, we leverage the strength of Bayesian methods (which yield full distributions over unknowns) to achieve point 3 above.

**Figure 1.**
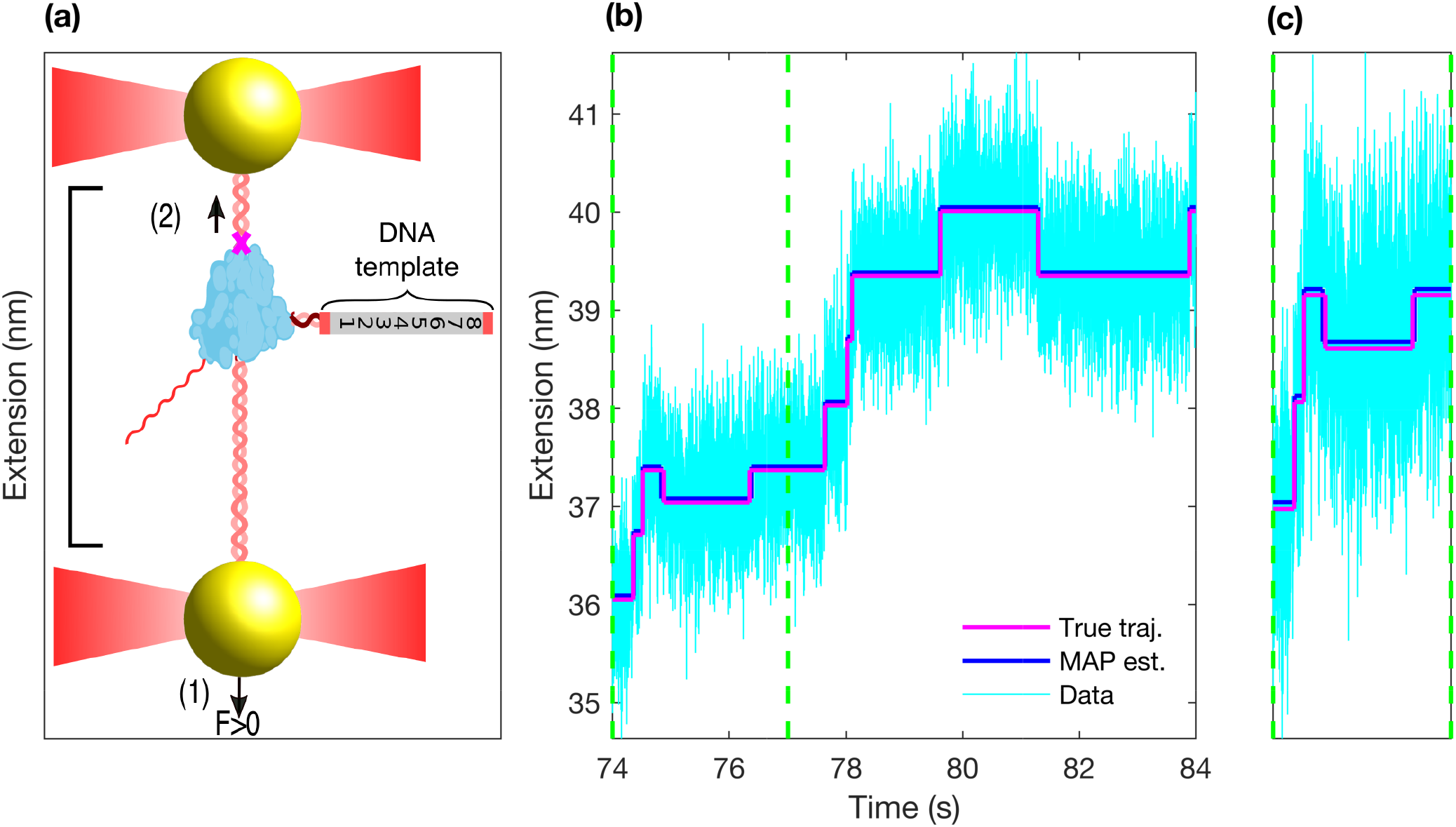
Schematic of a dual optical trap and the data generated. On panel (a), we have an illustration of an dual optical trap experiment. It shows RNAP actively transcribing a DNA template. Experimentally, in the dual optical trap setup, an assisting force (F > 0 in the arrow direction labeled with 1) may be applied to the bottom bead while the top bead is fixed in space. By “assisting” we mean that the force is applied in the same direction as the transcription of RNAP and producing an extension in the arrow direction labeled with 2. Opposing forces (where F < 0) are also possible. For each location change (translocation) of RNAP there is an extension measurement. This measurement gives rise to the signal on panel (b) that represents the extension between the two beads. Panel (b) contains a simulated extension time trace. In the panel (c) we are showing the regions defined with dashed green lines in panel (b). In panel (b) and (c) we also show the simulated true extension (in the absence of noise) between the optically trapped beads associated to RNAP’s progression (in magenta) as a function of time. The equivalent noisy extension as a function of time (with realistically added noise as described later in Model Description section is shown by the cyan. In this study our goal is, in part, to estimate the true extension (termed *maximum a posteriori,* or MAP, estimate) given this noise level under the added constraints that transition probabilities are location-dependent (i.e., heterogeneous in space).

## MATERIALS AND METHODS

In this study, we consider the conventional dual optical trap setup (20, 22, 24, 27). Precisely, in the experimental setup, a DNA template is attached to one optically trapped bead while RNAP is attached to the other bead (22, 24). Before the onset of the measurements, the two beads approach each other and subsequently RNAP attaches to the DNA template (22, 24). Upon this attachment, during the measurements, RNAP moves along the DNA template while it produces the transcribed RNA (20, 24). The motion of RNAP along the DNA template is represented in the measurements by the change of the distance between the beads (27). Depending on the force geometry employed in the experiment during the measurements, the measured extension may decrease (opposing force geometry) or increase (assisting force geometry) (24). The setup in Figure 1 illustrates the case when the experiment is carried out under the assisting force geometry. All experimental data here are described in (24). Below, we present our model and the method we developed for the analysis of the measurements.

### Model description

Here we describe the mathematical formulation of our method. We begin with the overall input that consists of the measured extension **x** = (*x*_1_,*x*_2_,*x*_3_,…,*x_N_*) where *x_n_* indicates the extension, associated with the translocation of RNAP, measured at equidistant time levels that we label *t_n_*, for *n* = 1,2,…,*N*; see Figures 1 and 2.

**Figure 2.**
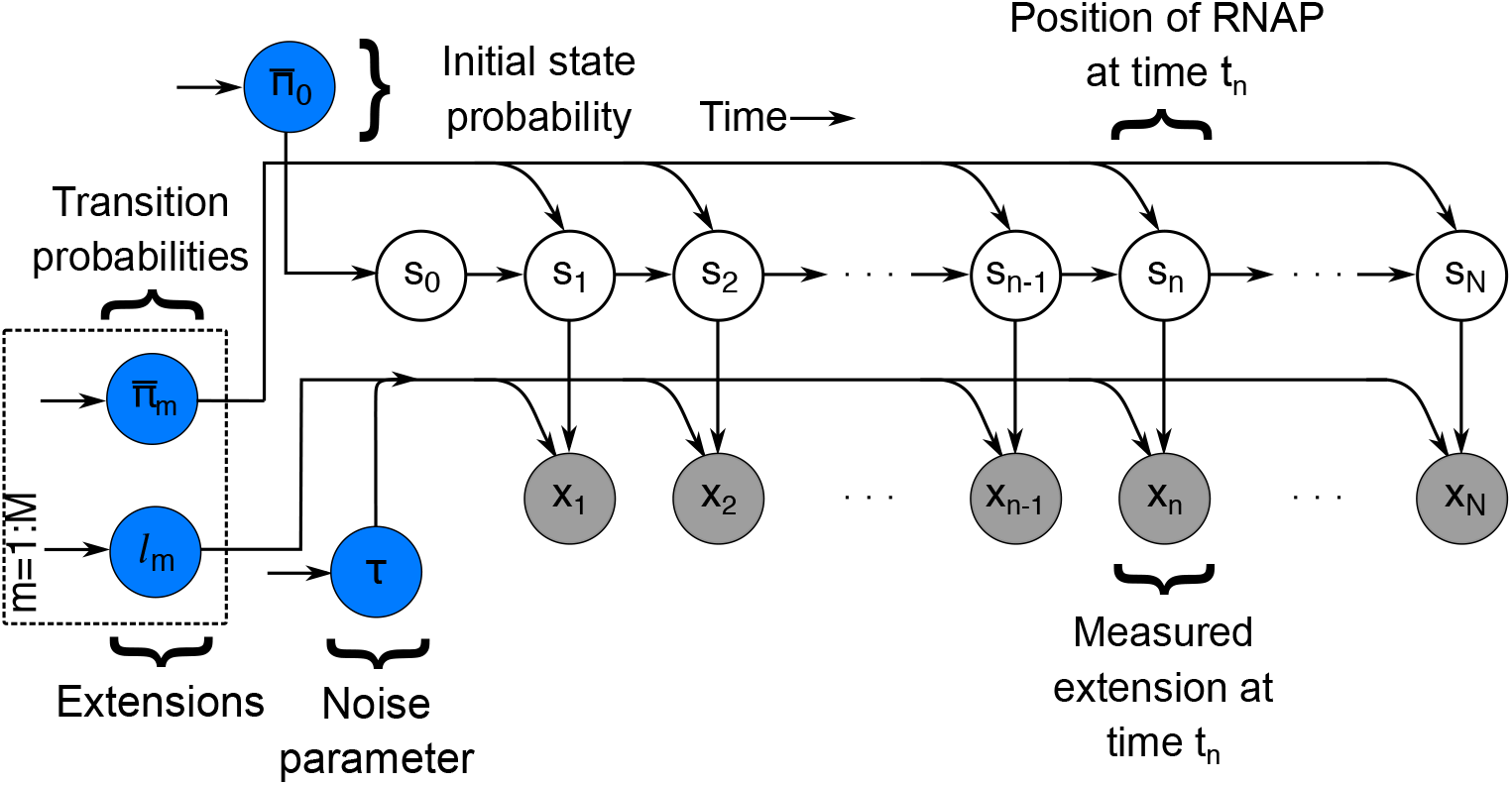
Graphical representation of the formulation used in the analysis of optical trap trajectories. By convention, quantities contained in gray circles denote observations and those contained in blue circles demand prior distributions. Here, a Markov chain *s_n_* denotes the location of RNAP on the transcribed DNA, the measurements are denoted by *x_n_*. Measurements are directly affected by the locations of RNAP namely *s_n_* at time *t_n_* and the measurement noise *τ* and extensions *l_m_* for all *n* = 0,1,…,*N* and *m* = 1,2,…,*M*. RNAP dynamics at each location *s_n_* during the time course of an experiment are governed by the transition probabilities *π_m_* for all *m* = 0,1,2,…,*M*.

We begin by defining *s_n_* which is the bp on the DNA template occupied at time level *t_n_.* Base pairs run from 1 (the begining bp) to the end of the transcribed region which is *M* (end bp). Naively, the average extension associated with this location is *s*_*n*_λ where λ is a conversion factor from nm to bp (0.33 nm/bp) (45).

As we avoid any pre-processing, the initial extension, xi onwards (typically the first few thousand of a total of tens to hundreds of thousands of data points), is set by the distance between the beads called the offset denoted by *l*\*. Thus the average *x_n_*, for any *n*, is *l*\* + *l_s_n__*.

In the presence of measurement noise, the recorded extension *x_n_* is stochastic and sampled from

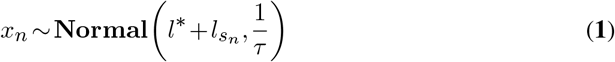

for *n* =1,2,…, *N*. Here *τ* is called precision and it has units of 1/nm^2^. Equation (**1**) should read as follows “*x_n_* is sampled from a normal (Gaussian) distribution with mean *l*\* + *l_s_n__* and variance 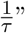.

In the presence of intrinsic noise (arising from the variability in bp jump size of RNAP depending on the bp DNA sequence and local environment), the extension, *l_s_n__* is also stochastic and sampled from

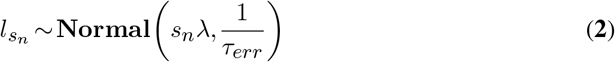

where 1 /*τ_err_* is again a variance with units of nm^2^.

Finally, the progression of RNAP from bp to bp, is governed by the Markovian assumption. Put differently, the probability of visiting the location *s_n_* (termed a “state” in the language of HMMs) at time level *t_n_* is determined exclusively by the state *s*_*n*–1_ occupied at the very previous time. In statistical notation, we write

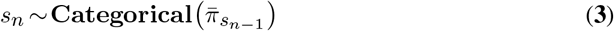

where 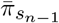 is the transition probability vector associated with transitions out of *s*_*n*–1_ including self-transitions (that is, the probability of staying put).

In this setting, when we speak of the transition probability vector associated with transitions out of the *m^th^* location, we write 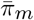. The transition probability matrix, denoted by 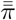, contains 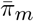 as its row and is a matrix of size *M* × *M* that encodes transitions from any location to any other location including self transition. The size *M* coincides with the total number of visited bp and by virtue of its value (as we have different matrix elements from neighboring bp to neighboring bp location) inherently imposes nonidentical transition matrix elements across all bps.

Armed with the data, **x**, and the generative model that incorporates both forward and backward steps, that we have just described, our goal now is to infer from the data those quantities that we care about. These include: RNAP locations *s_n_* for *n* =1,⋯,*N*, extensions *l_s_n__* for *n* = 1,⋯, *N* associated with locations *s_n_,* transition probabilities 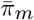 for *m* = 0, ⋯ *M* where 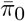 is the initial probability vector associated with RNAP’s 1^*st*^ location (namely *s*_0_).

### Model Inference

To determine full distributions over the quantities listed above, we follow the Bayesian paradigm. Other parameters not included in the above list (such as *τ_err_*, *l*\* and *λ*) can be also learned within the Bayesian paradigm. However, it is computationally more efficient to calibrate these parameters separately. For this reason, we set these parameters to known physical values within our framework. Details regarding the values of these parameters are provided in Supplementary Notes 1 and 4. We place prior distributions for the parameters 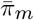,*s*_0_,*l_s_n__*, *τ* for all *n* = 1,2,…,*N* and *m* = 0,1,…,*M* that we wish to learn from the data. We discuss these priors below.

We start with 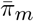 for all *m*=0,1,2,...,*M*. To estimate transition probabilities 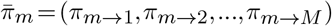 we place a sticky Dirichlet prior on each of 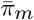. This prior is conjugate to the categorical distribution appearing in Equation (**3**) and it reads as follow

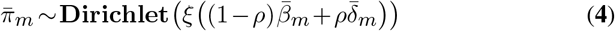

where 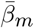 is a collection of hyperparameters that play the role of the base distribution (34), 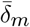 is the Dirac delta centered at *m*, *ξ* is known as the sticky parameter and *ρ* is the sticky proportion parameter. Larger *ρ* values promote self-transitions. Next we define the structure of the base distribution, 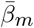.

We have two cases: the base distribution for the initial fictitious state *s*_0_ and that for all other states from 1,⋯, *M*. For the first case, as we can start from any state from 1 through *M*, the base distribution 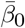 is

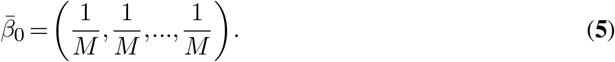

For the second case, only self-transitions, or steps one bp forward (for *m* =1,⋯, *M* – 1)or backward (for *m* = 2,⋯, *M*) within one time level to two steps forward (for *m* =1,⋯, *M* – 2) or backward (for *m* = 3,⋯, *M*) within one time level are allowed. The form for 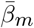 for *m* = 1, ⋯, *M* is specified in Supplementary Note 2.

Next we place a prior on *s*_0_ which is 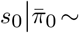 **Categorical** 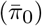. The evolution of the subsequent states is dictated by Equation (**3**). The prior on *l_s_n__* is dictated by Equation (**2**) and its hyperparameter *τ* is dictated by the conjugate prior 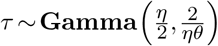, with hyperparameters *η* and *θ*. Details of the adopted distributions are provided in Supplementary Note 4. Once the choices for the priors given above are made, we form a joint posterior distribution (32–35, 46, 47)

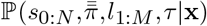

containing all unknown variables that we can learn. Due to lack of an analytical form for the posterior distribution, we developed a specialized computational scheme exploiting Markov Chain Monte Carlo (MCMC) methods to generate random samples from this distribution.

We explain the details of the implementation of our method in Supplementary Notes 3 and 4. A graphical summary of the entire formulation is shown in Figure 2.

## RESULTS

In this section, we first validate our method by computing posteriors over the unknown quantities 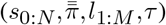 using simulated time traces, **x**, mimicking the properties of singlemolecule optical trap RNAP transcription experiments. We later move onto results from time traces from control *in vitro* experiments. The time required for the computations is provided in Supplementary Note 4.3.

### Demonstration and validation with simulated data

To demonstrate the robustness of our method, we simulate optical-trap time traces assumed to be collected at 800 Hz for 50 s. This is commensurate with the number of data points for the experiments we analyze. We simulate two cases: i) low measurement noise (high *τ*), see Figure 3; and ii) high measurement noise (low *τ*), see Figure 4. In Supplementary Figure 4, we provide the power spectral density plots of the analyzed time traces to show what the noise amplitudes are for both analyzed low and high measurement noise cases. The posteriors we obtain, in all our figures for the simulated data Figures 3 to 6 are informed from the analysis of only one time trace. To begin, in Figure 3 we simulate an optical trap transcription extension time trace with low measurement noise (1/*τ* = 0.25 nm^2^). The sample trace is shown in Figure 3 (a) where we zoom into a portion of panel (a) in panel (b) and panel (c). We first determine the posterior over the RNAP trajectories and, for illustrative purposes, show the *maximum a posteriori estimate* trajectory in Figure 3 (a), (b) and (c); Figure 5 shows the posterior over the average residence residence time distribution. The average residence time distribution in Figure 5 is obtained from the posterior over the transition probabilities and this is described in detail in Supplementary Note 4.

**Figure 3.**
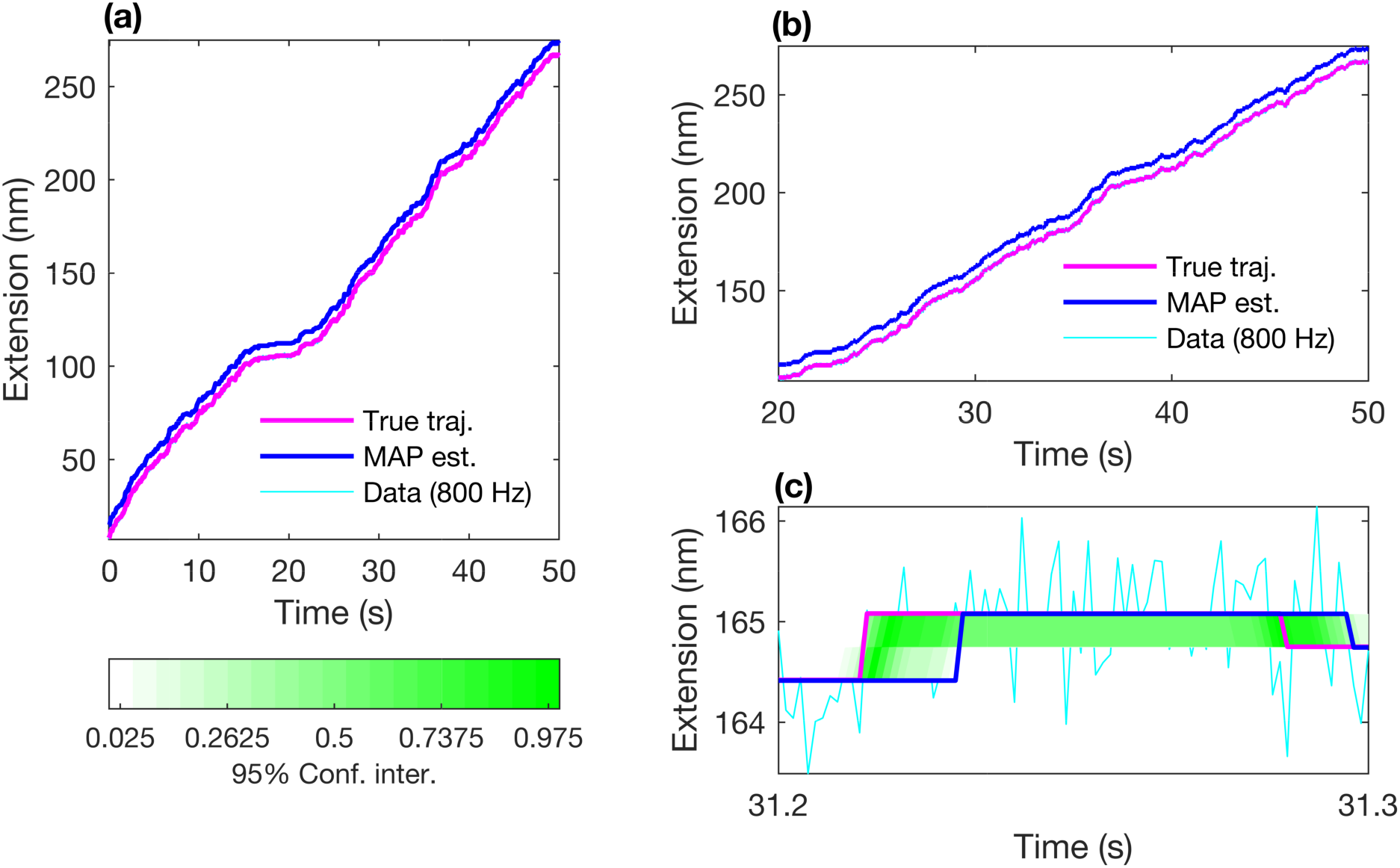
Simulated data analysis with low measurement noise 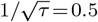 nm. In panel (a), simulated extension data in cyan is shown with parameters set to values provided in Supplementary Notes 2 and 4. As this is simulated data, the ground truth trajectory, *s*_1_,⋯,*s_N_*, is known and each *s_n_* corresponds to a bp on DNA. Therefore we know the associated trajectory with this ground truth trajectory by simply multiplying it with the λ = 0.33 nm/bp. In the generative model, we allow RNAP to take at most two steps at once. Therefore, in the trajectory we have some steps that are one or two times ~ 0.33 nm. This trajectory is given in panels (a), (b) and (c) by the magenta line. Using our framework, we determine the MAP estimate for the trajectory, blue line, that closely overlaps with the ground truth. For visual purposes we shifted the MAP estimate trajectory upward to avoid ambiguity in the comparison of trajectories. Panels (b) and (c) are zoomed in versions of panel (a). Further analysis for the measurement noise estimates is provided in Supplementary Figure 3.

**Figure 4.**
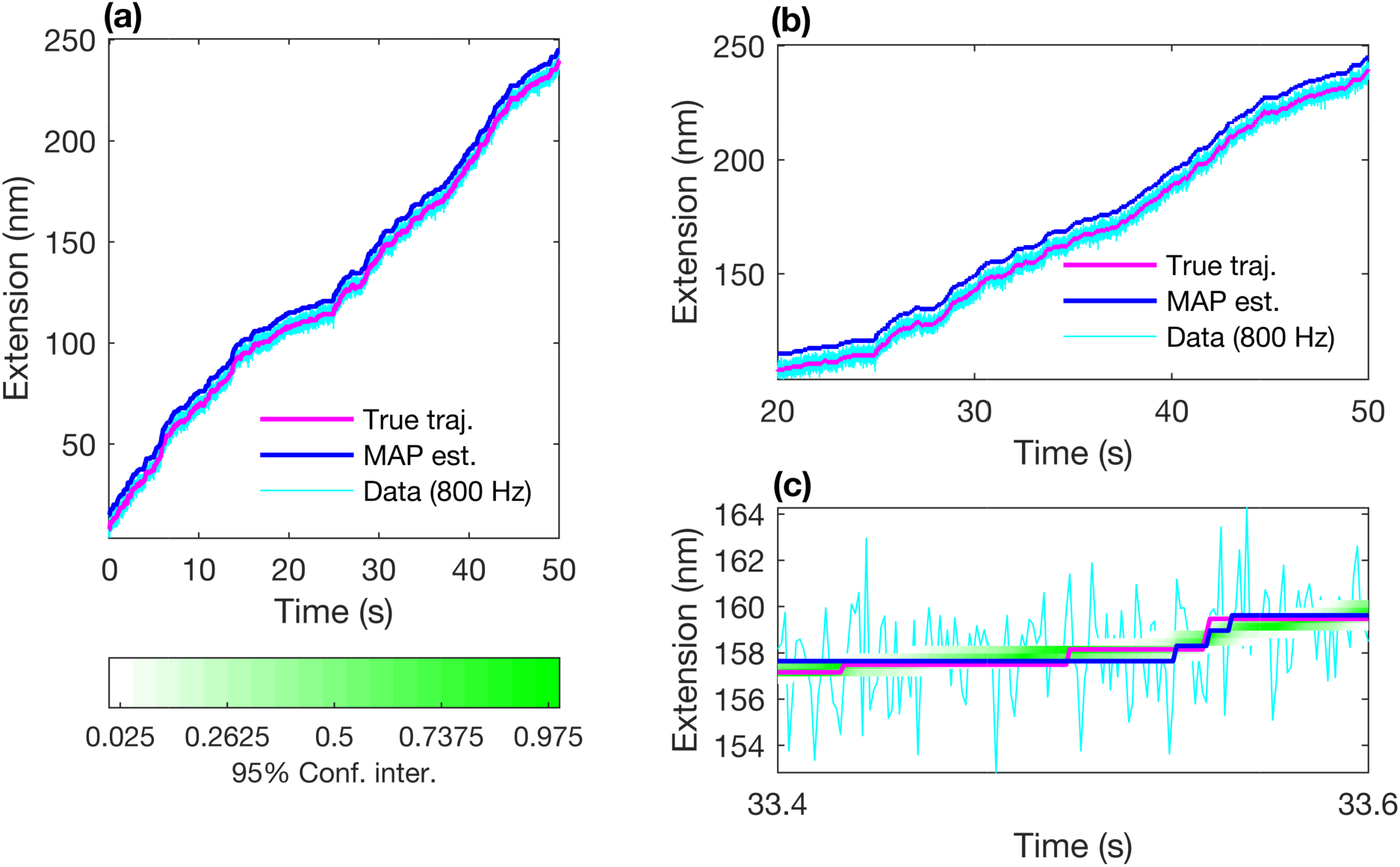
Simulated data analysis with high measurement noise 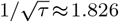 nm. As in Figure 3, we shifted the MAP estimate trajectory upward to avoid ambiguity in the comparison of the MAP estimate and true trajectories. Other than the value of *τ*, all other parameters are the same as in Figure 3. In panel (a), simulated extension data in cyan is shown with parameters set to values provided in Supplementary Notes 2 and 4. As this is simulated data, the ground truth trajectory, *s*_1_,⋯, *s_N_*. is known and each *s_n_* corresponds to a bp on DNA. Therefore we know the associated trajectory with this ground truth trajectory by simply multiplying it with the λ = 0.33 nm/bp. This trajectory is given in panels (a), (b) and (c) by the magenta line. Using our framework, we determine the MAP estimate for the trajectory, blue line, that closely overlaps with the ground truth. Panels (b) and (c) are zoomed in versions of panel (a). Further analysis for the measurement noise estimates is provided in Supplementary Figure 2.

**Figure 5.**
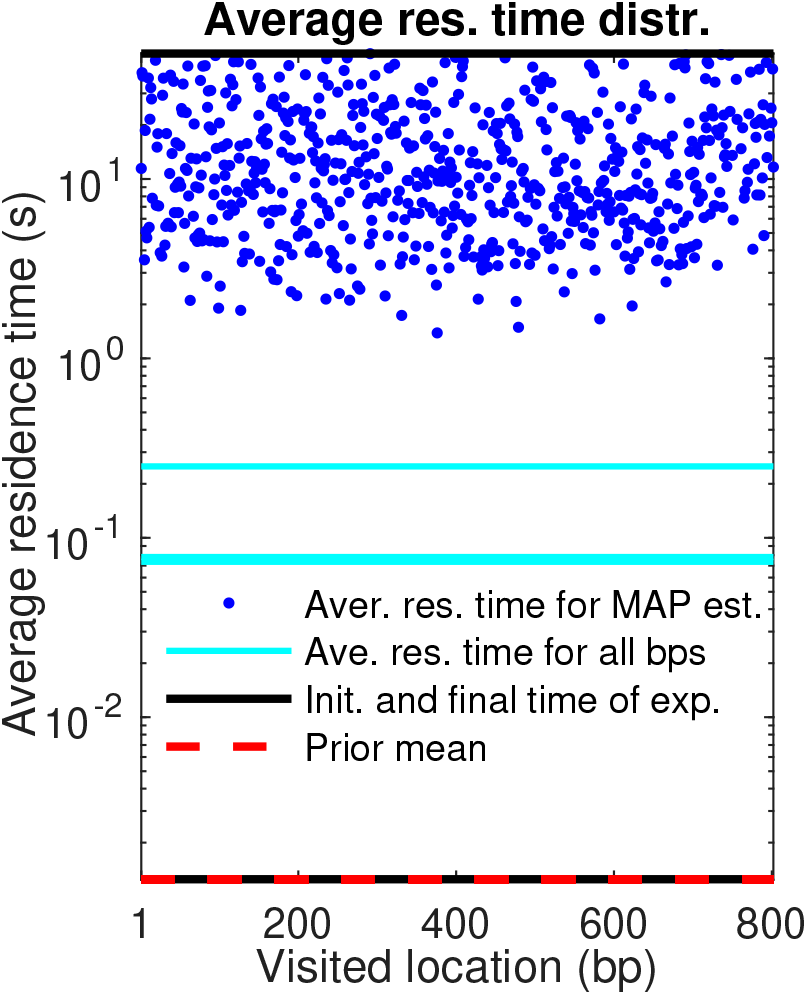
Analysis of average residence time distribution associated with the MAP estimate for the simulated data with our framework for the low measurement noise 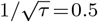 (nm). The minimum residence time is 0 and the maximum residence time is the end of the experiment. These two extremes are shown in black as the bottom most and top most lines, respectively. The ground truth for the average residence time is shown by the lines in cyan at (1/(1 – 0.983))Δ*t*,(1/(1 – 0.984))Δ*t*,(1/(1 – 0.995))Δ*t* where Δ*t* =1/800 s. We show the average residence time distribution for the MAP estimate with blue dots for each bp. Namely, we used the MAP estimate diagonal transition probabilities to calculate the average residence time for each bp as given in Equation (**6**). The reason behind the discrepancy between the exact value of the average residence times and its estimated distribution is due to the size of the unknown parameter space and limited data size. Although the model underestimates the exact distribution of the average residence times for every visited bp, the true trajectory is within the 95% confidence interval of the posterior distribution as we provided in Figure 3. All of our average residence time analysis relies on the residence times extracted from the MAP trajectories rather than the ones calculated from the transition probabilities. We provided the exact same analysis for the high measurement noise average residence time distribution associated with the MAP estimate in Supplementary Figure 4.

**Figure 6.**
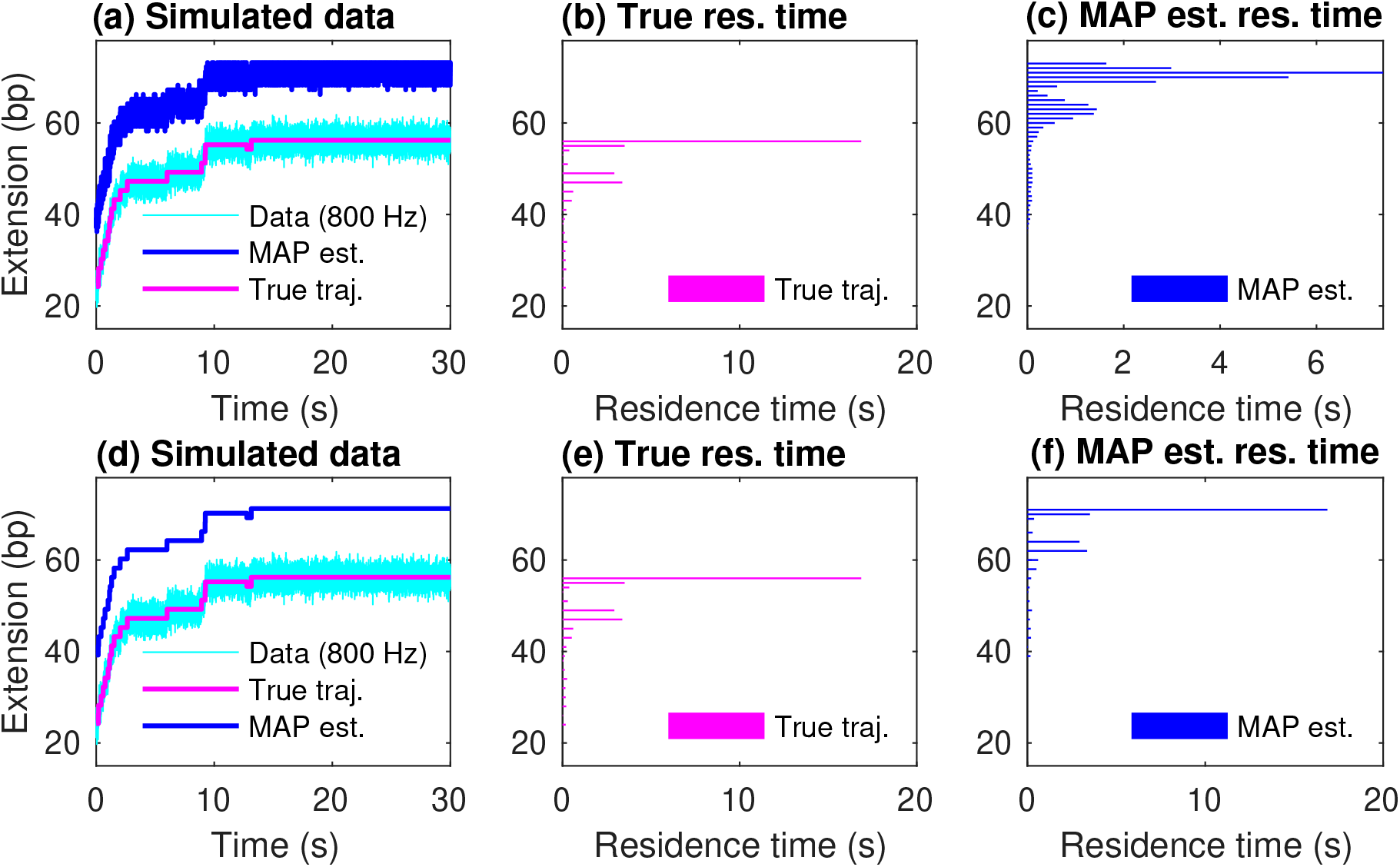
Analyzing the low noise regime data with a naive HMM and our framework. Here, simulated data is generated with measurement noise 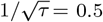 nm. Simulated data shown in panel (a) in cyan is generated with parameters set to values *M* = 49, *l*\* =8 nm for total 30 s measurement time with the transition probability matrix provided in Supplementary Note 2. As this is simulated data, the ground truth trajectory, *s*_1_,⋯, *s_N_* is known corresponding to the transcribed bps on DNA. Therefore the associated extension can be readily calculated by multiplying this sequence *s*_1_, ⋯, *s_N_* by 0.33 nm/bp. This extension is given by the magenta line. Using a plain HMM framework, we determine the MAP estimate for the trajectory, blue line, that overlaps poorly with the ground truth as it grossly underestimates the residence time from the noisy signal. As in Figures 3 and 4, we shifted the MAP estimate trajectories upward for visual clarity in the comparison of trajectories. In panel (c), we see the disagreement between the extracted residence time by the MAP estimates and ground truth residence time given in panel (b). We now analyze the same simulated data with our framework shown in panels (d), (e) and (f) where panel (e) provides the true residence information as in panel (b). In panel (d), we determine the MAP estimate for the trajectory given by the blue line, that overlaps very well with the ground truth. Panel (f) shows the true residence time histogram in magenta color and the residence time provided by the MAP estimate from our framework in blue color. We observe that the resulting MAP estimate from our framework performs quite well.

The breadth of the posterior (i.e., its variance) in Figures 3, 4 and 6 referenced above, reflects intrinsic noise levels such as thermal noise and the stochasticity in the elongation process. It also reflects the fact that the data is finite. In other words, if we had more data, the posterior would be sharpened but only up to a point. It retains some width on account of the stochasticity inherent to transcriptional elongation.

We briefly discuss in greater depth the MAP trajectory estimate, Figures 3 and 4 (a), (b) and (c). From Figure 3 (a), we see how closely the MAP estimate and the true simulated trajectory are overlapping. These MAP estimates can be used to extract residence time. We observe from the simulated data analysis, MAP estimates provided by standard HMM performs poorly in approximating residence time associated with every bp as shown in panel (c) of Figure 6. Instead, our method performs quite well in learning the correct residence time given in panel (f) of Figure 6.

As we have access to the full posterior of trajectories, we have also knowledge of the 95% confidence interval associated with every step in that path. The confidence interval is determined by the lower and upper limits of 95 percentile of the sampled trajectories. This confidence interval is shown as the green-white color curve of Figures 3 and 4 (c).

Next, we discuss the average residence time distribution in Figure 5. Due to the Markovian dynamics for the location transitions, the average residence time distribution associated with every *m^th^* bp denoted by *“T_m_”* is a function of self-transition probabilities, *π_m→m_*, and the time between time levels, Δ*t*. That is,

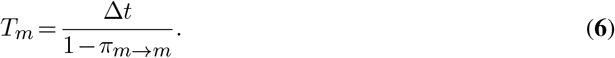

Detailed information regarding the derivation of the formulae for the average residence time distribution is provided in Supplementary Note 4 and the notations are listed in Supplementary Table 1.

### Experimental data

Next, we apply our method to experimental time traces obtained from dual ultra-stable optical traps (24). Within our framework, we analyze individual time traces under different experimental force geometries (assisting, opposing) as well as in the absence and presence of different elongation factors (GreB). Here, we consider 6 experimental setups. These are: 1) assisting force geometry with no addition of elongation factors (Figures 7 and 10 to 12); 2) assisting force geometry in the presence of GreB (Figures 10 to 12); 3) assisting force geometry in the presence of RNaseA (Figures 10 to 12); 4) opposing force geometry with no addition of elongation factors (Figures 8 and 10 to 12); 4) opposing force geometry in the presence of GreB (Figures 10 to 12); 6) opposing force geometry in the presence of RNaseA (Figures 10 to 12). In this section we focus on the 1^*st*^, 2^*nd*^, 4^*th*^ and 5^*th*^ experimental setups.

**Figure 7.**
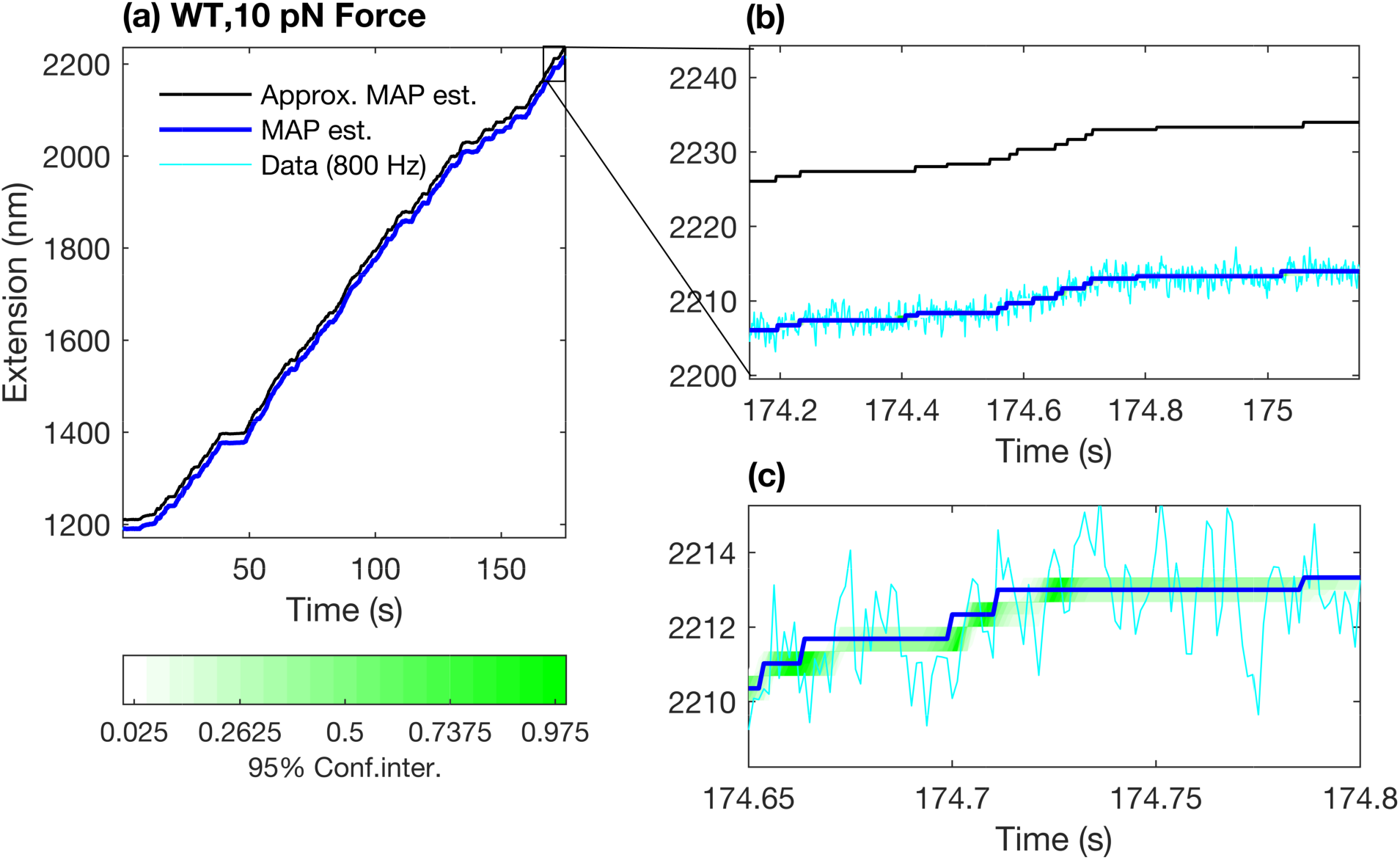
Analysis of +10 pN experimental trace with our framework in the absence of GreB and RNaseA. In panel (b) of this figure we zoomed in a small region of one of the analyzed traces for +10 pN shown in panel (a) and we superpose the MAP trajectory as well as the approximate MAP trajectory on the experimental trace. Here the approximate MAP trajectory stands for the trajectory that is closest to the MAP trajectory with respect to *l*^2^ – norm. Further zoomed experimental trace, MAP trajectory and the approximate MAP trajectory are provided in panel (c). Details of this approximate MAP trajectory is provided in Supplementary Note 3. As it was mentioned earlier in Figures 3 and 4, here we shifted the approximate MAP trajectory upward for visual reasons.

**Figure 8.**
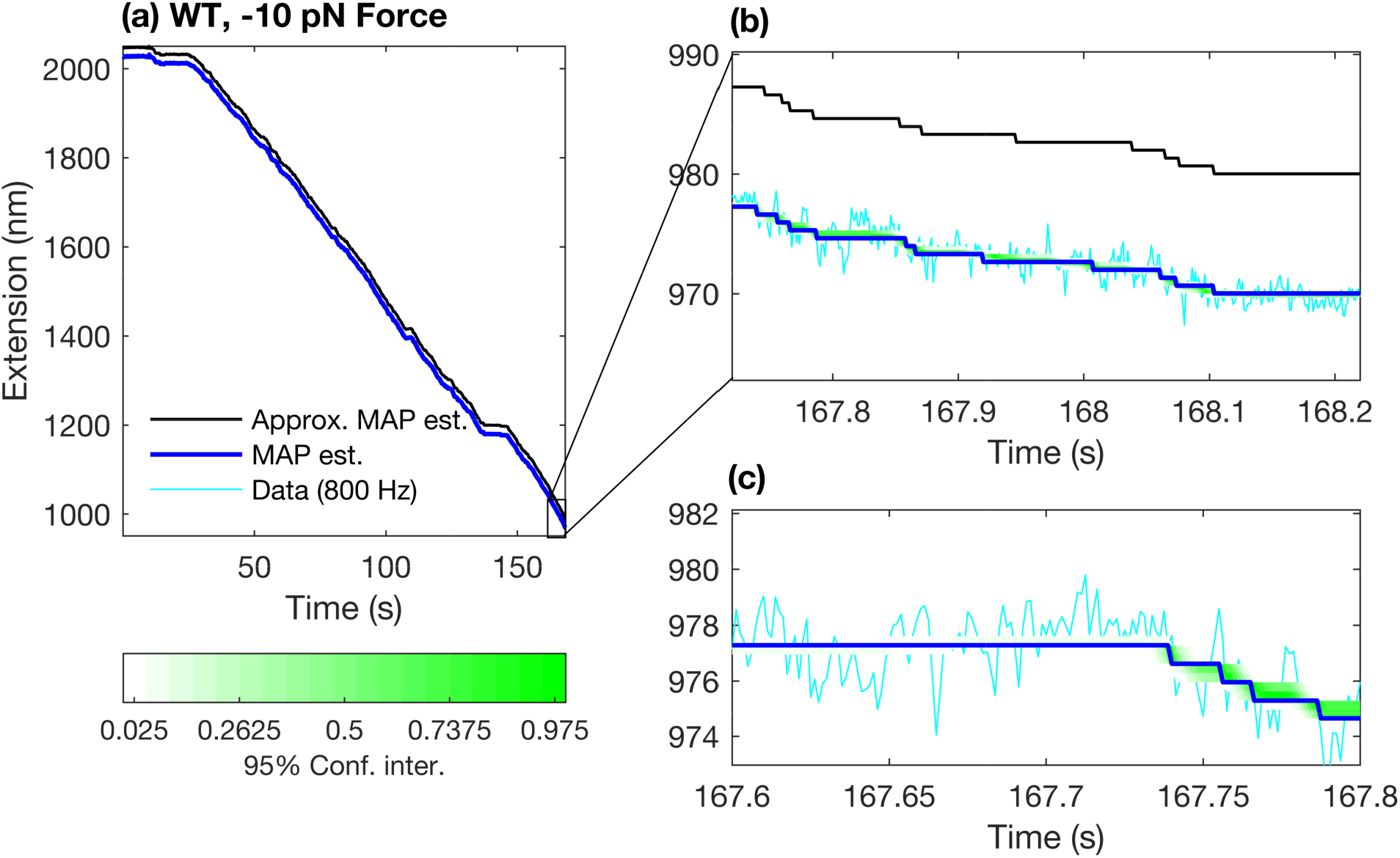
Analysis of −10 pN experimental trace with our framework in the absence of GreB and RNaseA. Here, we show an analyzed trace under −10 pN force. Other than the analyzed experimental time trace all the explanation about the panels in the figure is the same as in Figure 7. The approximate MAP trajectory is shifted upward for visual reasons.

A brief note on the types of DNA being transcribed is noteworthy here. These DNA contain 8 repeats each with identical sequence. However the length of nascent RNA is different for each of these repeats due to the experimental conditions including assisting and opposing geometries as well as the addition of RNaseA and elongation factor GreB (24). Next, the role of GreB is to rescue the backtracked RNAP (48) while RNaseA is responsible for cleaving the nascent chain (25).

### There is no clear distinction between short and long residence time

In Figure 7 (a) and Figure 8 (a), we show samples of analyzed experimental time traces associated with +10 pN and −10 pN forces. These help illustrate how the transitions between the steps of the MAP trajectory and approximate MAP trajectory look like with forces of the same magnitude but opposing directions. We present the definition of the approximate MAP trajectory in Supplementary Note 3.

While the MAP trajectory is shown in Figure 7 (a) and Figure 8 (a), within the Bayesian framework, we obtain the full posterior distribution over all variables in the model and in Figure 7 (b) and Figure 8 (b) we provide the 95% confidence intervals over the trajectories.

This posterior distribution allows us to obtain distributions for residence times. To interpret the terminology provided by the existing literature on the classification of the residence times (20, 22, 24), in Figure 9, we superpose average residence time histograms for 10 trajectories (obtained from one experimental trace by resampling the posterior) and corresponding double and single exponential fits to these histograms.

**Figure 9.**
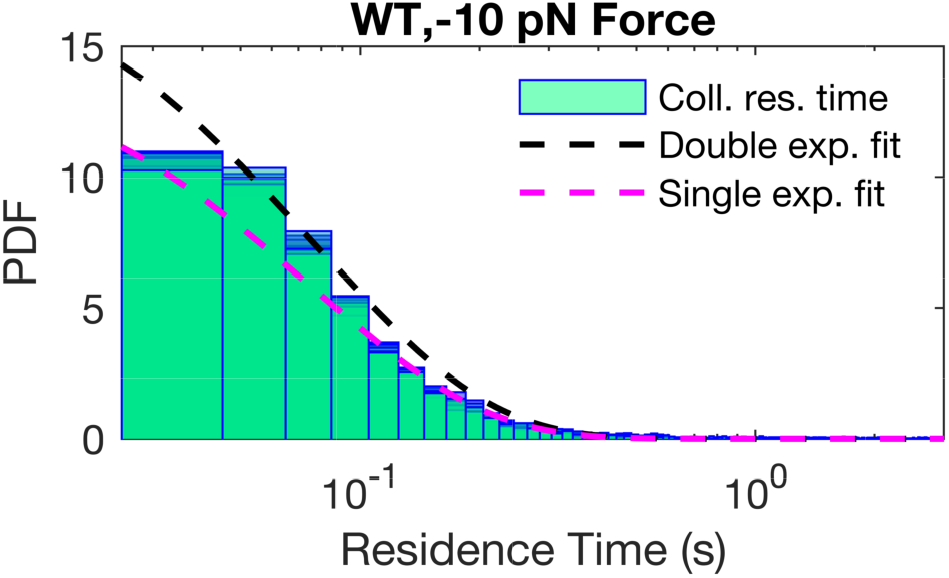
Double exponential dwell time distributions fitted to multiple sampled trajectories. Here we show the analysis regarding multiple sampled trajectories besides the MAP estimate with our tool for one experimental trace under −10 pN force in the absence of GreB or RNaseA. We combined the residence times over all repeat regions for each randomly chosen 10 trajectory. This type of analysis of the residence times for each experimental time trace is what we call as *“collective residence time”* abbreviated as “coll. res. time”. In this figure, we show how the PDFs of collective residence time for each trajectory are distributed. We should note that, double exponential fits as good as the single exponential to these residence time PDFs. Therefore, there is no clear distinction between short and long residence times for the RNAP transcription dynamics based on how residence time PDFs are fit to exponential distributions.

We should note that the quality of the exponential fits depends on how representative the chosen sampled trajectories are. We showed the analyzed experimental time trace in Figure 8. In Figure 8 panel (b), we zoomed in the analyzed time trace and provided the 95% confidence interval along with the MAP estimate. The sharp width of the posterior (as suggested by the 95% confidence interval) suggests that the majority of trajectories closely resemble the MAP estimate. As can we sample multiple trajectories from the posterior, perhaps surprisingly, we find that some histograms over residence times are better fit by single and others by double exponentials. This suggests that variations around the MAP estimate may raise questions on the validity of single versus multi-exponential fits (20–22). Next, we report our results for the comparison of first 2 experimental setups along with the 4^*th*^ and 5^*th*^ experimental setups.

### Average residence times are not appreciably affected by the presence of GreB

We carried out our analysis for the experimental time traces associated with RNAP transcription in the presence of anti-backtracking (48) or elongation factor (24) GreB (0.87*μ*M) (24). We compare these results to the control experimental setups where the elongation factor GreB is absent. In our analysis, effects of GreB on transcriptional residence time are investigated based on 3 quantities (1) average residence time; 2) average backtracking duration; and 3) average backtracking counts. All these quantities were analyzed in two ways: 1) evaluating them over the entire 8 DNA tandem repeat regions and 2) evaluating them over the 2-4, 4-6 and 6-8 repeat regions. The reason for this is to determine the net effect of the length of the nascent chain on each of the 3 quantities of interest.

We find that there is limited effect of GreB on the average residence time over each of the DNA tandem repeat regions or the repeat sections as we show in Figure 10 (b) when we compare it with the control experiment where GreB is absent, that is Figure 10 (a).

**Figure 10.**
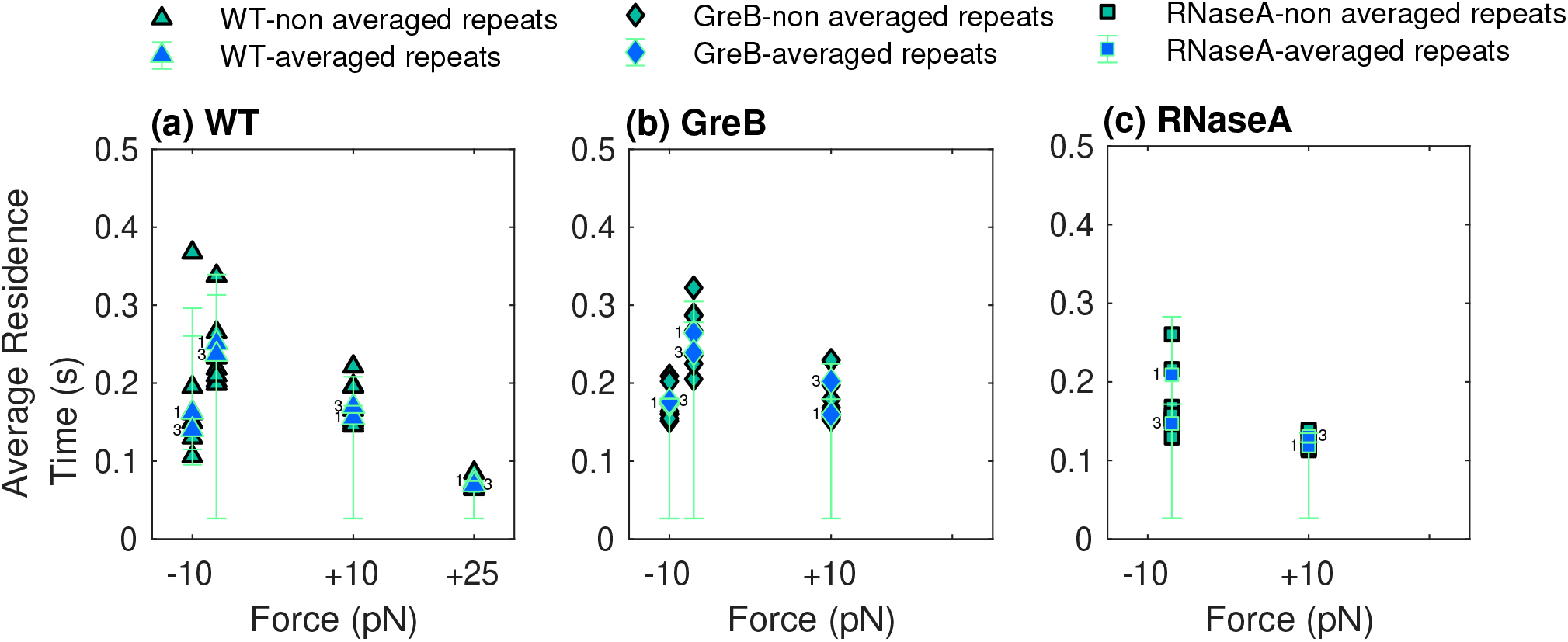
Average residence time for the collective analyses of experimental traces under various experimental conditions. Here we focus on the average residence time point statistic under different experimental conditions. Average residence time is calculated separately for all experimental conditions. In this manuscript, we analyzed 5 experimental traces for +25 pN force geometry, 5 traces for +10 pN, 3 traces for −10 pN and 5 traces for −7 pN. In addition, in the presence of GreB, 8 experimental traces under +10 pN, 6 traces for −10 pN and 7 traces for −7 pN force geometries were analyzed. Besides, in the presence of RNaseA, we analyzed 5 experimental traces for +10 pN and 5 traces for −7 pN force geometry. We produced samples between 4000 − 13000 for each analysis. There are 8 repeat regions for each analyzed trace. We emphasize that the analysis of the average residence time is not for *single nucleotides* but for repeat regions that are engineered with 239 bps. We superposed the information for the same repeat regions for the analyzed multiple traces from the same experimental conditions. The term “average” comes from the average of the superposed residence time information coming from multiple analyzed time traces for each repeat region. We extracted the residence time information from the MAP estimate trajectory for each analyzed experimental time trace. In panel (a), we show the analysis of the average residence time for RNAP transcription in the absence of elongation factors. Each green triangle corresponds to one of the 8 repeated region. In order to assess whether we observe different residence time across different repeats, where the sequence of translated RNA is the same but the length of the nascent chain is different, we averaged repeats 2,3,4 and represented this average by the blue triangle that we labeled 1. We did the same for repeats 4,5,6 that we labeled with the blue triangle labeled 2 and 6,7,8 with a blue triangle labeled 3. We repeated this analysis in the presence of GreB in panel (b) and RNaseA in panel (c) at concentrations of 0.87μM and 0.1mg/ml, respectively. We used 75% confidence interval obtained from all calculated residence time for each repeat section. These confidence intervals are shown by green lines superposed on the point estimates associated with the repeat sections. The main results of the figures above are discussed in Results.

The study in (20) investigates the effects of GreB on pausing (short-long) events. These events are defined based on their duration and this duration starts from the cessation of the transcription elongation until RNAP recovers from it and starts forward translocating. In (20), it is reported that GreB was effective in reducing long pausing events that last longer than their identified threshold 20 s. As we do not pickup significant differences between short and long residence time, we therefore do not note an effect of GreB at the 75% confidence interval from panel (b) of Figure 10. Our result regarding the limited effects of GreB is also consistent with (22, 49).

In the next section we focus on 1^*st*^, 3^*rd*^, 4^*th*^ and 6^*th*^ experimental setups to investigate the effects of RNaseA on average residence time.

### The effect of applied forces is abolished in the presence of RNaseA

We further investigate the presence of RNaseA (0.1 mg/ml) (24) on the average residence time dynamics of RNAP with our framework.

Here, our objective is to investigate the relation between the nascent chain and the average residence time dynamics of RNAP. Thereby, we analyze experimental traces in the presence and absence of RNaseA, namely in the absence and presence of the nascent chain, respectively.

When we analyze the data with our framework, we find that RNaseA reduces the variance in the average residence time distribution over the repeat regions under the opposing applied force under opposing force geometry as shown in Figure 10 (c).

As the absence of nascent chain (due to the presence of RNaseA) leaded to the reduced effects of the opposing force geometry on the averaged residence time of RNAP, we make an analogy of “shoe-lace effect” for this contribution of the nascent RNA over the RNAP residence time dynamics.

In the literature, there has been discussions on the effects of nascent RNA on the residence time dynamics of RNAP through its interaction with RNAP prior to RNAP’s translocation (24) yet it remains unproven. Moreover, in (25, 30, 40), researchers investigated the effects of transcribed DNA’s chemical composition on residence time of RNAP based on the interaction of the nascent chain with RNAP. They found that upon removal of the nascent chain, the chemical composition effects of the transcribed DNA template on residence time dynamics of RNAP disappears. Yet these results rely on heavy data pre-processing to eliminate detected backtracks as well as defining the short and long residence times prior to the analysis (40).

Next, we would like to investigate the role of GreB and the nascent chain (therefore the absence or presence of RNaseA) on the backtracking motion of RNAP.

To begin, in (22, 24, 48), researchers mentioned that GreB is responsible for rescuing RNAP upon RNAP’s backtracking as little as 2 bp. Additionally, in (24), they did not find a direct communication between the role of nascent chain and backtracking dynamics of RNAP from their analysis. Inspired from these studies, here, we analyze the experimental time traces in the presence of GreB (24) with our framework to further investigate GreB’s effects on backtracking motion of RNAP based on the average backtracking duration and counts.

We analyze the experimental traces (24) in the presence of RNaseA to investigate the interplay between the backtracking motion of RNAP and the role of nascent chain.

Thereby, in the next section, we look into all the experimental setups as we listed previously especially we will investigate the roles of RNaseA and GreB from the perspective of two point estimates including average backtracking duration and backtracking counts.

### Average backtracking durations show little change in the presence of GreB

In this section, we probed the effect of RNaseA (0.1 mg/ml) (24) and GreB (0.87*μ*M) (24) on the average backtracking durations and counts.

First, we start with the analysis of the effect of RNaseA on the average backtracking durations under both assisting and opposing force geometries. Our results indicate that, as shown in Figure 11, there is slight change in the average backtracking durations in the presence and absence of RNaseA over the repeat sections at the 75% confidence interval.

**Figure 11.**
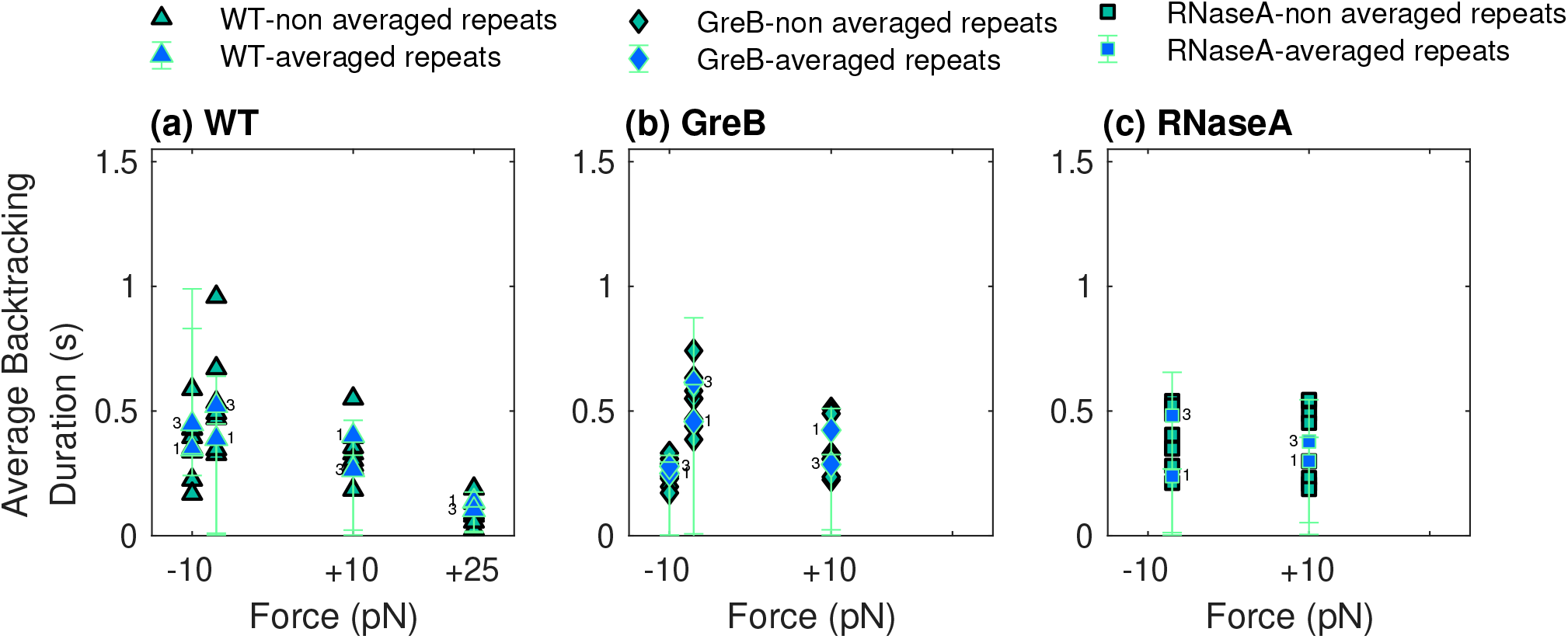
Average backtracking durations obtained by collective analyses of experimental traces under various experimental conditions. Here, we obtained the average backtracking durations from the experimental time traces given as follows. We analyzed 5 experimental traces for +25 pN force geometry, 5 traces for +10 pN, 3 traces for −10 pN and 5 traces for −7 pN. In addition, in the presence of GreB, in panel (b), 8 experimental traces under +10 pN, 6 traces for −10 pN and 7 traces for −7 pN force geometries were analyzed. Besides, in the presence of RNaseA, in panel (c), we analyzed 5 experimental traces for +10 pN and 5 traces for −7 pN force geometry. We calculated the backtracking durations for each repeat region from the MAP estimate trajectory for all analyzed experimental time traces separately. Then we superpose the backtracking duration information for the repeat regions and then we found the average backtracking duration from this new list of backtracking durations. We repeated this process for all experimental conditions. In panel (a) we show the analysis of the mean backtrack durations associated with each repeat region for the wild type RNAP transcription. Each triangle corresponds to a repeat region. We later combined the mean of backtrack durations along the repeats 2,3,4 and label it as 1 and 6,7,8 as the new repeat section 3 and investigated the backtrack duration distribution of these repeat sections. We repeated this analysis in the presence of GreB in panel (b) and RNaseA in panel (c) at concentrations of 0.87μM and 0.1mg/ml, respectively. We used 75% confidence interval obtained from all calculated backtracking durations for each repeat section. This confidence intervals are shown in green lines super imposed on the point estimates associated with the repeat sections.

As in (24), our results suggest that this slight effect of RNaseA namely, removal of the force dependence of backtracking duration, is stronger for the opposing force geometry case. This might be due to relatively lower transcription rates of RNAP under the opposing force geometry (22, 24). Therefore, there would be enough time for the secondary structures to form from the nascent RNA (24, 25). This may be the reason of the nascent RNA’s enhanced effect on the residence time of RNAP during its transcription elongation under opposing force.

Next, we investigated how GreB affects backtracking motion of RNAP by analyzing first, its effects on the average backtracking duration and average backtracking counts.

Figure 11 shows that there is limited effect of GreB, namely reduced average backtacking durations with 75% confidence.

This slight effect of GreB on the average backtracking duration can be attributed to the less than sufficiently long residence times recovered with our framework. In (20, 50), it is mentioned that GreB rescues the RNAP from a location after it spends long time (longer than 20 s) at that location. In our framework, we have not observed residence times longer than 20 s over all of the force geometries. We believe that this might be the reason behind our observation regarding the limited effect of GreB on the averege backtracking durations. These findings are consistent with (20, 51).

Finally, we analyze how backtracking counts are correlating with the applied forces (Figure 12) in the presence of RNaseA and transcription elongation factor GreB. Our analysis provides support on account of force dependence for backtracking counts in experimental traces with no RNaseA or GreB. Namely, it is expected to observe more backtracking counts under opposing force geometry than the assisting force geometry as explained in (50).

**Figure 12.**
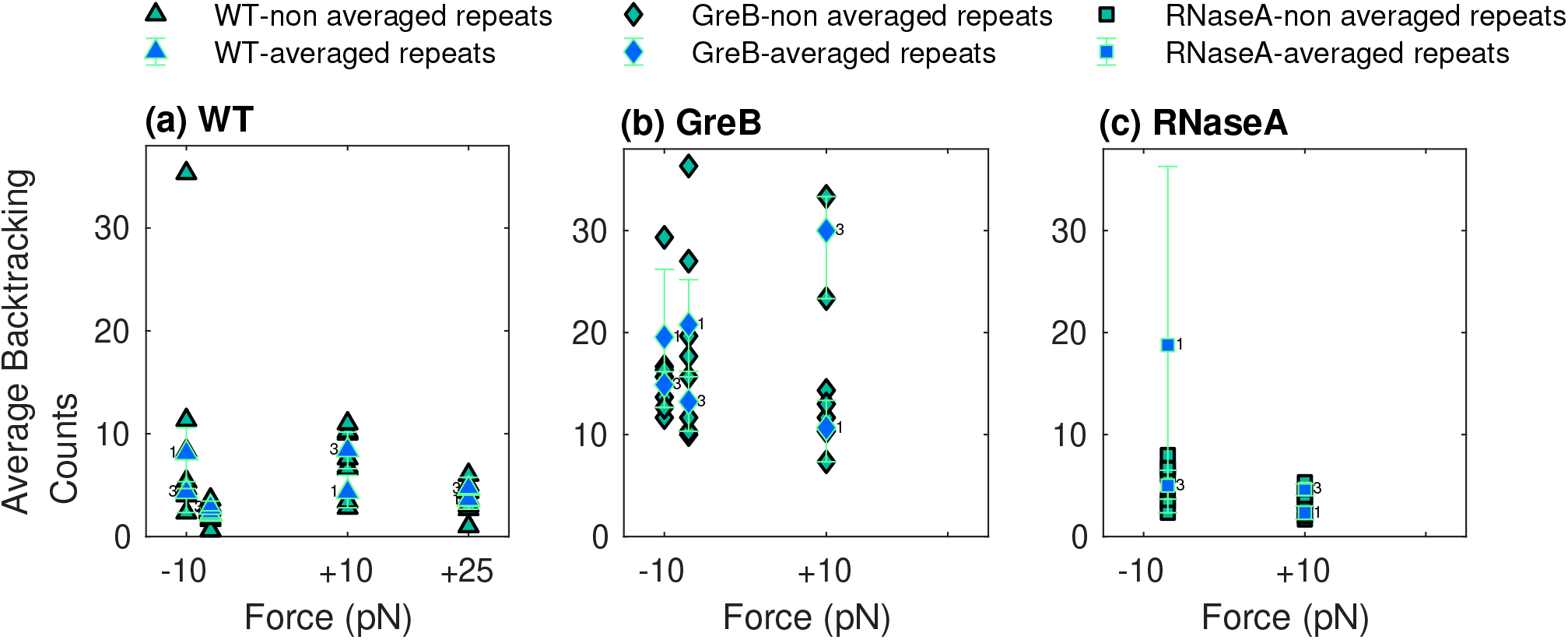
Average backtracking counts through the collective analysis of experimental traces under various experimental conditions. Our analysis provided in this figure relies on the analysis of the following traces. We analyzed 5 experimental traces for +25 pN force geometry, 5 traces for +10 pN, 3 traces for −10 pN and 5 traces for −7 pN. In addition, in the presence of GreB, in 8 experimental traces under +10 pN, 6 traces for −10 pN and 7 traces for −7 pN force geometries were analyzed. Besides, in the presence of RNaseA, we analyzed 5 experimental traces for +10 pN and 5 traces for −7 pN force geometry. We calculated the backtracking counts for each repeat region from the MAP estimate trajectory for all analyzed experimental time traces separately. Then we superpose the backtracking count information for the repeat regions and then we found the average backtracking counts (here average is calculated with respect to the total number of analyzed time traces) from this new list of backtracking counts. We repeated this process for all experimental conditions. In panel (a), we show the analysis of average backtrack counts for the wild type RNAP transcription. Each triangle corresponds to a repeated region and later we averaged backtrack counts along the repeats 2,3,4 and label it as 1 and 6,7,8 as the new repeat section 3 and investigated the backtrack count distribution of these repeat sections. We repeated this analysis in the presence of GreB in panel (b) and RNaseA in panel (c) at concentrations of 0.87μM and 0.1mg/ml, respectively.

## DISCUSSION

In this study, we developed a Bayesian approach to use Markov models to systematically investigate transcription elongation dynamics of *E. Coli* RNAP by analyzing experimental time traces obtained from dual optical trap assays. Our method has a number of unique features. First, unlike other kinetic models (16, 52–54), our method does not assume that transcription elongation dynamics of RNAP consists of well separated short or long residence time events. In addition, by contrast to earlier studies, we also allow the transcriptional dynamics to vary based on the visited locations by RNAP on the DNA template via location-dependent transition probabilities. In our analysis, we treat extensions *l_m_*, for *m* =1,2,…,*M* stochastically and assume these are sampled from a distribution that depends on the detected location *s_n_* for *n* = 1,2,…,*N*. As such, *l_m_* is treated as a variable and learned in our Bayesian framework.

Typically, the way in which previous studies get good statistics and error bars is by collecting many experimental time traces (41, 42). Here, on the other hand within error bars (as propagated from data uncertainty within the Bayesian paradigm), we find a number of models all which could be consistent with the data. The latter is true even as we increase the amount of data analyze. In other words, this suggests that within error many pause models could potentially be consistent with data.

### No discrimination between short and long residence times

We analyzed multiple experimental time traces under a variety of applied forces with and without GreB and RNaseA. We provide the list of analyzed time traces for every experimental condition in Supplementary Figures 6-15. We demonstrated the applied force dependence of various point statistics with our framework including average residence times, average backtrack durations and average backtrack counts over the repeats even when we have the average with respect to the corresponding DNA sequence of the analyzed traces. We also demonstrated the average backtracking lengths in Supplementary Figure 15. Contrary to existing literature, we emphasize that we do not find posterior probabilities for residence times appreciably longer than 1 s though we can still infer small posterior probabilities for finding residence times that are longer than 1 s. This is demonstrated from the analysis illustrated in Figures 9 and 10.

### Shoe-Lace effect of nascent RNA

Ultimately, we find that the effect of the elongation factor GreB on the average residence time is quite limited. In the past, the effect of GreB was investigated on the long pauses where a pause means the interval starting with the cessation of the elongation that lasts until the RNAP recovers from it and starts forward translocating (20). Specifically, in (20), they reported that GreB reduces the duration of long pauses that last longer than 20 s. We speculate that because our framework did not recover probable trajectories with long residence times, as long as 20 s, we could not see GreB affecting the duration of such undetected events.

On the other hand, in the presence of RNaseA (namely when the nascent RNA is removed), the variance in the average residence time distribution over the repeat regions under the opposing applied force is reduced. We described this finding with our analogy of the “shoe-lace effect” of nascent RNA on the residence time dynamics of RNAP during transcription.

Previously, researchers in (22, 24, 25) found that the nascent chain modulates the residence time of RNAP. In (40), it was shown that the removal of the nascent RNA abolishes the effects of the chemical composition of the transcribed DNA on the residence time dynamics. Therefore our result, regarding the nascent RNA’s effect on the residence time dynamics of RNAP during transcription, aligns with the existing literature (22, 24, 25, 40).

### Presence of GreB has limited effects on average backtracking durations

Now we turn to backtracking. The reason is, as mentioned in (20, 22, 24, 30, 50), that backtracking can be considered as a process that plays the role of proofreading in RNAP transcription in addition to providing a mechanism for transcription termination (50). Although it is believed that the long residence times are providing enough time for the transcription activities to be regulated (24, 25, 30), backtracking plays a role in helping elongation factor (GreB) to enforce RNAP for controlling its accuracy in transcription and translocation (50).

Interestingly, what we find from our backtracking analysis is that there is limited effect of GreB on average backtracking durations over the repeat regions that is more apparent under −10 pN force than the assisting forces with 75% confidence. Indeed, backtracking is known to be favored under the opposing force geometry in the literature (50). Therefore, GreB facilitates protruding RNA 3’-end cleavage and thus rescues RNAP from longer backtracking durations (24, 50, 55). This leads RNA 3’-end to align well with the RNAP active site again. Hence, we observe reduced average backtracking durations as soon as GreB acts on RNAP during its transcription elongation.

Similarly, based on our analysis of the experimental time traces in the presence and absence of RNaseA, nascent RNA did not appear to have any effect in backtracking durations. However, the average number of backtracking counts turned out to be reduced in the absence of the nascent RNA. This suggests that, as the nascent RNA gets longer it is more likely for the nascent RNA to interact with the RNAP before it translocates (24). This might give rise to higher number of backtracks for RNAP to manage its accuracy in transcription (50). Therefore the average number of backtracks become higher in the presence of the nascent RNA.

## CONCLUSION

We have used an approach based on the Bayesian sticky HMMs to interpret time traces obtained from dual optical trapping experiments associated with RNAP’s transcription elongation. Our approach enabled us to analyze these time traces for RNAP’s transcription elongation without denoising them. Upon the analysis of time traces with our approach, we estimated the trajectories of RNAP with the MAP trajectory estimate. We used MAP trajectory estimates to learn RNAP’s transcription elongation dynamics. In doing so, we calculated various point statistics from the MAP trajectory estimate including: average residence time, average backtracking duration and average backtracking counts. According to the analysis of these point statistics, our approach revealed the following: 1) there is no clear separation between short and long residence times; 2) the presence of RNaseA reduces the variance of average residence time distribution. Therefore, this result suggests that nascent RNA affects the average residence time dynamics of RNAP. We term the effect of the nascent RNA on RNAP the “shoe-lace effect”; 3) the presence of GreB has limited effects on the average residence time and average backtracking duration.

## SUPPLEMENTARY INFORMATION

Supplementary information, source code and GUI versions (see Supplementary Note 1 and Supplementary Figure 1) of the methods developed are provided seperately.

## AUTHOR CONTRIBUTIONS

Z.K. developed computational tools and analyzed data; I.S. contributed computational tools; Z.K., I.S. and S.P. conceived the research; S.P. oversaw all aspects of the project.

## ACKNOWLEDGEMENTS

SP acknowledges support from NSF CAREER grant MCB-1719537 and NIH NIGMS (R01GM134426). ASU cluster AGAVE and Saguaro are the main computational resources utilized in this study. We thank Carlos Bustamante, Ronen Gabizon and Antony Lee for their helpful suggestions in orienting the initial thoughts on this project. ZK thanks Ufuk Kilic, M. Kemâl Ozalp, Laurie M. Straube for their helpful suggestions on the manuscript.

